# Differential stoichiometry among core ribosomal proteins

**DOI:** 10.1101/005553

**Authors:** Nikolai Slavov, Stefan Semrau, Edoardo Airoldi, Bogdan Budnik, Alexander van Oudenaarden

**Affiliations:** Department of Bioengineering, Northeastern University, Boston, MA 02115, USA; Department of Statistics and FAS Center for Systems Biology, Harvard University, Cambridge, MA 02138, USA; Leiden Institute of Physics, Leiden University, 2333 CC, Leiden, The Netherlands; Hubrecht Institute, Royal Netherlands Academy of Arts and Sciences and University Medical Center Utrecht, Uppsalalaan 8, 3584 CT, Utrecht, The Netherlands

## Abstract

Understanding the regulation and structure of ribosomes is essential to understanding protein synthesis and its dysregulation in disease. While ribosomes are believed to have a fixed stoichiometry among their core ribosomal proteins (RPs), some experiments suggest a more variable composition. Testing such variability requires direct and precise quantification of RPs. We used mass-spectrometry to directly quantify RPs across monosomes and polysomes of mouse embryonic stem cells (ESC) and budding yeast. Our data show that the stoichiometry among core RPs in wild-type yeast cells and ESC depends both on the growth conditions and on the number of ribosomes bound per mRNA. Furthermore, we find that the fitness of cells with a deleted RP-gene is inversely proportional to the enrichment of the corresponding RP in polysomes. Together, our findings support the existence of ribosomes with distinct protein composition and physiological function.

## Introduction

Ribosomes catalyze protein synthesis but have only a few characterized roles in regulating translation (Mauro and Edelman, 2002; Xue and Barna, 2012). Rather, the most–studied molecular regulatory mechanisms of translation are mediated by eukaryotic initiation factors, RNA binding proteins, and microRNAs (Hendrickson *et al*, 2009; Fabian and Sonenberg, 2012). The characterized catalytic role of the ribosomes corresponds well to the model of the ribosome as a single complex with a fixed stoichiometry: 4 ribosomal RNAs and 80 core RPs (Warner, 1999; Ben-Shem *et al*, 2011), some of which are represented by several paralogous RPs. Despite the longstanding interest in ribosome structure and function, the exact stoichiometry and possible heterogeneity of the ribosomes have been challenging to measure directly (Weber, 1972; Westermann *et al*, 1976; Hardy, 1975). Such measurements are enabled by modern quantitative mass spectrometry (MS). Indeed, MS has transformed our understanding of protein complexes, such as proteasomes (Wang *et al*, 2007) and nuclear pore complexes (Ori *et al*, 2013), by demonstrating variability among their protein subunits. Furthermore, quantitative MS has proved useful in characterizing ribosome biogenesis (Chen and Williamson, 2013).

Studies of eukaryotic ribosomes (Mazumder *et al*, 2003; Galkin *et al*, 2007; Komili *et al*, 2007; Kondrashov *et al*, 2011; Horos *et al*, 2012; Lee *et al*, 2013) have demonstrated that (*i*) genetic perturbations to the core RPs specifically affect the translation of some mRNAs and not others and (*ii*) mRNAs coding for core RPs are transcribed, spliced, and translated differentially across physiological conditions (Ramagopal and Ennis, 1981; Ramagopal, 1990; Parenteau *et al*, 2011; Slavov and Dawson, 2009; Slavov and Botstein, 2011, 2013; O’Leary *et al*, 2013; Slavov *et al*, 2014; Gupta and Warner, 2014; Jovanovic *et al*, 2015). These results suggest the hypothesis (Mauro and Edelman, 2002; Gilbert, 2011; Xue and Barna, 2012) that, depending on the tissue type and the physiological conditions, cells can alter the stoichiometry among the core RPs comprising the ribosomes and thus in turn alter the translational efficiency of distinct mRNAs. Alternatively, differential RP-expression can reflect extra ribosomal functions of the RPs (Mazumder *et al*, 2003; Wool, 1996; Warner and McIntosh, 2009). Furthermore, polysomes (multiple ribosomes per mRNA) from different cancer cell-lines have similar core RP stoichiometries (Reschke *et al*, 2013). Thus, the variable RP stoichiometry in the ribosomes of wild-type cells that is suggested by the ribosome specialization hypothesis remains unproven.

We sought to test whether wild-type cells have ribosomes with differential RP stoichiometry. For this test, we chose two divergent eukaryotes: budding yeast *Saccharomyces cerevisiae*, and mouse ESC. We chose budding yeast because of our previous observations that RPs are differentially transcribed across growth-rates (Slavov and Botstein, 2011, 2013) and that RP levels change differentially between glucose and ethanol carbon source (Slavov *et al*, 2014). To investigate whether such differential transcription of RPs affects the ribosomal composition, we used the same media as in our previous experiments, minimal media supplemented with 0.2 % glucose. In this media, unlike in rich media supplemented with 2 % glucose, yeast cells have a prominent monosomal peak that may reflect different translational regulation (Ashe *et al*, 2000; Castelli *et al*, 2011; Vaidyanathan *et al*, 2014). We chose embryonic stem cells to test differential RP stoichiometry in wild-type mammalian cells because of the interesting phenotypes of RP deletions/knockdowns in ESC. For example, haploinsufficiency for Rps5, Rps14, or Rps28 interferes with ESC differentiation but not with their self-renewal (Fortier *et al*, 2015). Furthermore, unlike heteroploid cancer cell-lines grown in culture, ESC have a high monosomes-to-polysomes ratio, consistent with the possibility of differential translational regulation (Sampath *et al*, 2008; Fortier *et al*, 2015).

## Results

### Differential stoichiometry among core RPs in mouse ESC

To explore whether the stoichiometry among core RPs can vary, we first isolated monosomes and polysomes from exponentially growing mouse embryonic stem cells (ESC), doubling every 9 hours, Figure S1A. The ESC ribosomes were isolated by velocity–sedimentation in sucrose– gradients (Figure 1A); see Methods. To confirm that the prominent monosomal peak is reflective of ESC biology and not of poor ribosome fractionation, we also fractionated the ribosomes of neuroprogenitor cells derived from the ESC. Despite growing three times slower (doubling time 29 hours) than the ESC, the neuroprogenitor cells have a larger fraction of their ribosomes in polysomal complexes, Figure S1B. This observation confirms earlier findings by Sampath *et al* (2008), and thus further bolsters the conclusion that a low polysome-to-monosomes ratio is characteristic of ESC.

**Figure 1.**
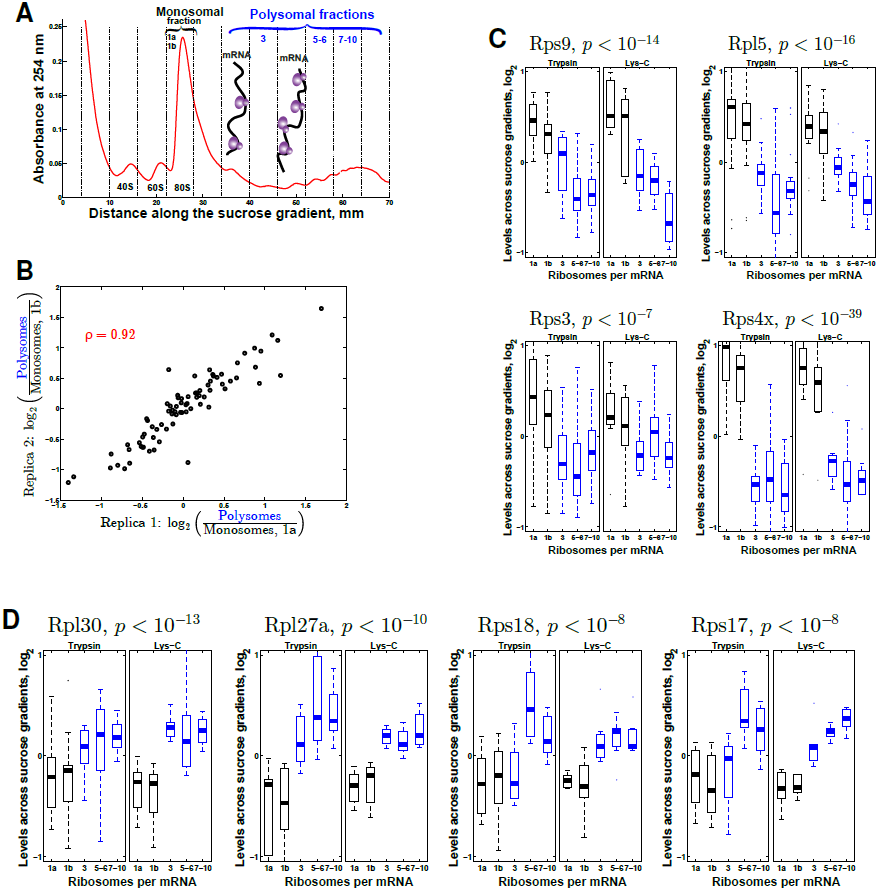
The stoichiometry among core RPs in mouse ribosomes depends on the number of ribosomes per mRNA. (A) Sucrose gradients allow separating ribosomes that are free or bound to a single mRNA (monosomes, depicted in black) from multiple ribosomes bound to a single mRNA (polysomes, depicted in blue). The absorbance at 254 nm reflects RNA levels, mostly ribosomal RNA. The vertical dashed lines indicate the boundaries of the collected fractions. Fractions are labeled at the top with numbers reflecting the number of ribosomes per mRNA. (B) Replicates MS measurements of the monosomes (1a and 1b) indicate reproducible estimates for RP enrichment in polysomes. (C, D) Some RPs are enriched in monosomes (C) and others in polysomes (D). The relative levels of each RP are quantified as the median levels of its unique peptides, and the probability that the RP levels do not change across the quantified fractions is computed from ANOVA (indicated at the top). The distributions of levels of all unique peptides from trypsin (left panels) and from lys-C (right panels) digestions are juxtaposed as boxplots to depict the consistency of the estimates across proteases, different peptides, and experiments. For each fraction, the mean intensity of all RP peptides was normalized to 1. On each box, the central line is the median, the edges of the box are the 25*^th^* and 75*^th^* percentiles, and the whiskers extend to the most extreme data points. See also Figure S1 and Figure S2.

Having isolated monosomes and polysomes, we sought to quantify their protein composition. The proteins from individual sucrose fractions were digested to peptides, labeled with tandem mass tags (TMT), and quantified on Orbitrap Elite based on the MS2 intensities of the TMT reporter ions; see Supplemental Information. The monosomal sample was quantified in two replicates (1a and 1b), and the results indicate very high reproducibility (*ρ* = 0.92; Figure 1B). To control for protease and peptide biases, the proteins from each analyzed sucrose fraction were digested either by trypsin (T) or by lys–C (L) and peptides from each digestion quantified independently. Because of the different specificity of trypsin and lys-C, most RP peptides (1058) were identified and quantified only in the trypsin or only in the lys–C digestion, while only 269 peptides were identified and quantified in both digestions. Thus, only very few peptide-specific biases (such as co-isolation interference) may be shared between the two digestions.

The measured levels of a unique peptide (a peptide present in a single RP) reflect the levels of the corresponding RP, post–translational modifications (PTMs) of the peptide (if any), and measurement error. We quantify on average ten distinct RP–peptides per RP (Figure S2A), and the levels of these peptides allow both the estimation of the RP levels and the consistency of these estimates. To depict both the estimates and their consistency, we display the full distributions of relative levels of all peptides unique to an RP as boxplots in Figure 1C, D. The RP levels across the sucrose gradient (estimated as the median of the levels of unique peptides) indicate that some RPs are enriched in monosomes (Figure 1C), while other RPs are enriched in polysomes (Figure 1D). Each RP group includes proteins from both the large (60S) and the small (40S) subunits of the ribosomes and thus differential loss of 40S or 60S cannot account for the RP levels displayed in Figure 1C, D. Indeed, normalizing for the total amount of 40S and 60S proteins in each fraction does not alter significantly the results. The RP enrichment in Figure 1 is substantially higher than the measurement noise, consistent across replicates and across distinct peptides, and highly statistically significant at false discovery rate (FDR) < 10*^−6^*. The relative levels of all RPs with quantified unique peptides are displayed in Figure 2 to illustrate the global pattern of RP levels across monosomes and polysomes. This pattern shows more RPs whose variability is consistent across replicates and enzymatic digestions. In contrast, the levels of RPs buried in the core of the ribosomes remain constant, with estimates fluctuating within the tight bounds of the measurement noise, Figure 2. This fixed stoichiometry among RPs constituting the ribosomal core suggests that even ribosomes lacking some surface RPs likely have the same core structure.

**Figure 2.**
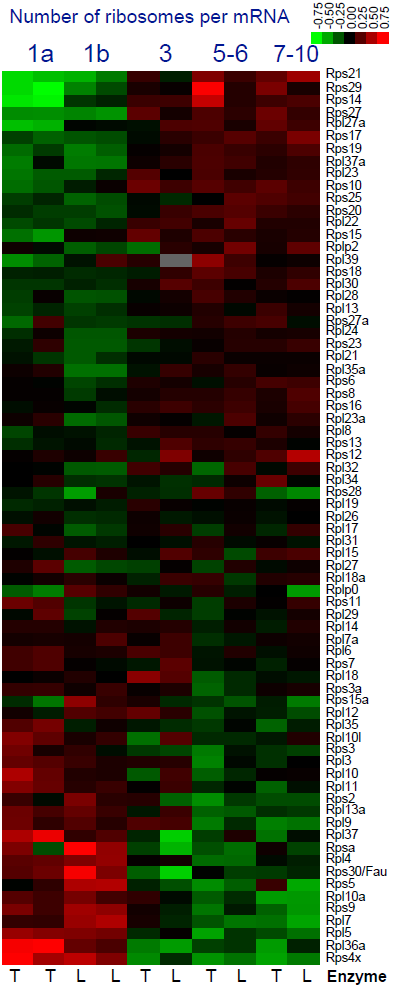
Global pattern of differential stoichiometry among mouse RPs across sucrose gradients. Global pattern of differential stoichiometry among mouse RPs across sucrose gradients. The relative levels of core RPs in monosomes and polysomes were quantified by MS and found to vary depending on the number of ribosomes bound per mRNA. The measurement noise was estimated by (*i*) replica quantification of the monosomal fraction (by using different tandem-mass-tags reporter ions, 126 or 131) and by (*ii*) estimating RP levels separately using either trypsin (T) or lys-C (L) digestion, as indicated at the bottom of each column. The log_2_ levels of each RP are shown relative to their mean. See also Figure S2, Figure S3 and Tables S1-S3.

In principle, if only a few peptides are quantified per RP, the measured peptide variability might reflect reciprocal variability in corresponding PTM isoforms (if any) across the sucrose gradients, e.g., the unmodified isoform is enriched in monosomes and a phosphorylated isoform is enriched in polysomes. Such differential distribution of PTM isoforms (if any) is interesting since it represents another layer of ribosome regulation, but cannot explain the data for an RP quantified by dozens of peptides spanning the protein length and indicating highly–consistent fold–changes across the sucrose gradient; see Figure 1, Figure S2, and Supplemental Information.

We further tested the differential RP stoichiometry with an independent method, Western blots, and in another strain of mouse ESC. Consistent with the MS data in Figure 2, the Western Blots data (Figure S3) indicate that Rps29 and Rps14 are enriched in polysomes, Rpl11 is enriched in monosomes, and Rpl32 does not change beyond the measurement noise.

### Differential stoichiometry among core RPs in yeast

Having found differential stoichiometry among mouse RPs, we sought to further explore (*i*) whether such ribosome heterogeneity is conserved to budding yeast and (*ii*) whether the RP stoichiometry can change with growth conditions and metabolic state. To this end, we employed sucrose gradients to separate the ribosomes from yeast cells grown in minimal media with either glucose or ethanol as the sole source of carbon and energy (Slavov *et al*, 2014); see Supplemental Information. Consistent with previous observations that the type and the concentration of the carbon source influence the ratio of monosomes to polysomes (Ashe *et al*, 2000; Castelli *et al*, 2011; Vaidyanathan *et al*, 2014), the ratio of monosomes to polysomes in our yeast cells grown in 0.4 % ethanol (Figure 3A) or in 0.2 % glucose (Figure 3B) is higher than is typically observed for yeast grown in rich media containing 2% glucose. As in mouse, some RPs are enriched in monosomes (Figure 3C) and others in polysomes (Figure 3D, E). This enrichment is reproducible (correlation between replicates *ρ* = 0.97; Figure 3F) and consistent across independent unique peptides whose levels are shown as boxplot distribution in Figure 3C-D.

**Figure 3.**
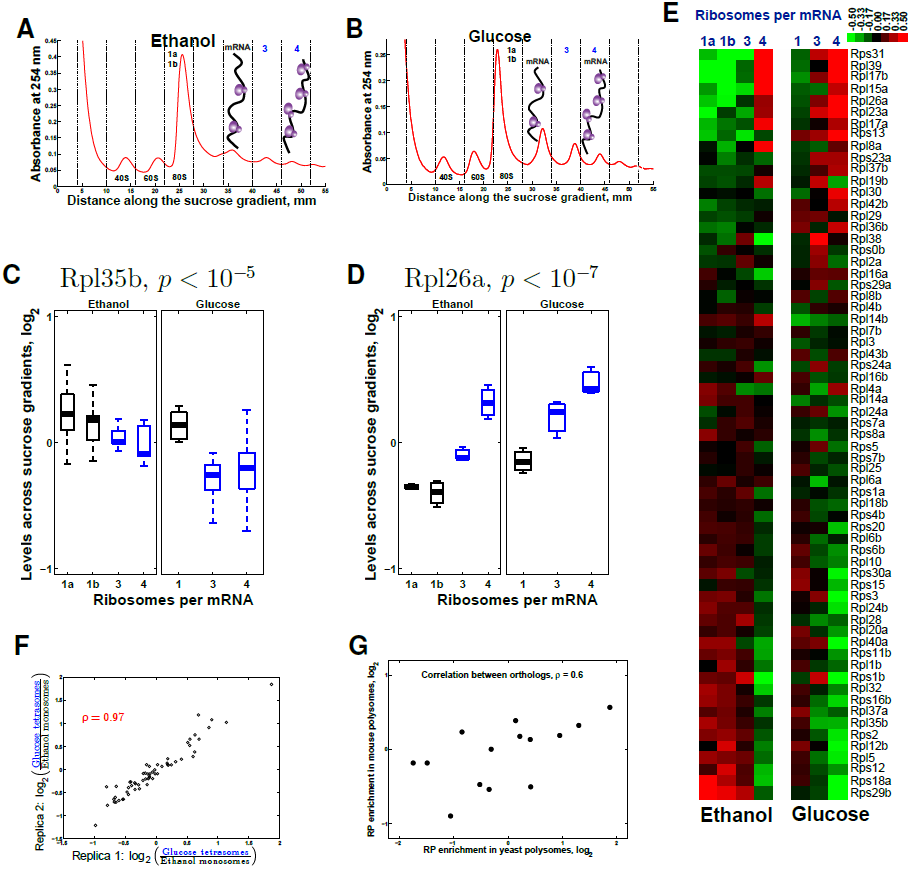
The stoichiometry among core RPs in yeast ribosomes depends both on the number of ribosomes per mRNA and on the physiological condition. (A, B) Ribosomes from either ethanol (A) or glucose (B) grown yeast were separated by velocity sedimentation in sucrose gradients. Depiction is as in A. (C) Rpl35b is enriched in monosomes (*p* < 10*^−3^*) and in ethanol carbon source (*p* < 10*^−3^*). Depiction is as in Figure 1. The p value at the top is computed from ANOVA and quantifies the probability of observing the variability of Rpl35b peptides by chance. (D) Rpl26a is enriched in polysomes (p < 10*^−9^*) and in glucose carbon source (p < 10*^−4^*). (E) Levels of core RPs in the sucrose fractions estimated from their unique peptides quantified by MS. The RP levels vary depending on the carbon source (glucose or ethanol) and on the number of ribosomes bound per mRNA, indicated at the top. Monosomes from ethanol grown yeast were quantified in two biological replicates (first two columns). The log_2_ levels of each RP are shown relative to their mean. See Supporting Movie 1 and PDB files for color-coded depiction of these data on the 3D structure of the yeast ribosome. (F) The RP fold–changes between the tetrasomes of yeast grown in glucose carbon source and the monosomes of yeast grown in ethanol carbon source are highly reproducible. The ethanol samples were collected and processed independently and compared to the glucose tetrasomes. (G) The log_2_ ratios between polysomal and monosomal levels of mouse RPs are plotted against the corresponding log_2_ ratios of their orthologous yeast RPs. The significant (p-value < 0.03) positive correlation between these ratios suggests that the differential RP stoichiometry is conserved across yeast and mouse. The plot includes all orthologous RP pairs with over 65% sequence identity between yeast and mouse. See also Figure S2, Figure S4 and Tables S4-S5.

We investigated whether the differential levels of RPs, both in yeast and in mouse, may reflect the presence of ribosome biogenesis complexes or other extra-ribosomal complexes containing RPs. We estimated that biogenesis factors are over 200-fold less abundant than RPs across all samples (Figure S4A), and 80-fold less abundant even in the monosomal fractions (Figure S4B) where ribosome biogenesis particles are enriched; see Supplemental Information. These data suggest that the proteins derived from immature ribosomes can contribute about 1 − 3% to the RP fold-changes, while some measured RP fold-changes exceed 100 % (Figure 1). The contribution of immature ribosomes to our RP estimates can be further tested by using the order in which RPs are incorporated into the small subunits. This order has been established for bacterial RPs in vitro (Mulder *et al*, 2010) and confirmed in vivo (Chen and Williamson, 2013). We used this order, as well as the correspondence and nomenclature between orthologous bacterial and mammalian RPs (Jenner *et al*, 2012), to test the trends that are expected if biogenesis particles are abundant enough to influence RP quantification: RPs that are incorporated early should be enriched in the monosomal fractions and depleted from polysomal fractions; the late RPs should show the converse trend. While these trends are observed for some RPs (such as S4 and S14), the opposite trends are observed for other RPs (such as S3, S5, S11, and S15), Figure S4C. The overall pattern of relative RP levels in Figure 2 cannot be fully accounted for by the order of RP incorporation during ribosome biogenesis (Figure S4C).

The pattern of relative RP levels shown in Figure 3C-E indicates that RP stoichiometry depends on two factors: on the number of ribosomes per mRNA (as in mouse) and on the carbon source in the growth media; the RP levels that are higher in glucose compared to ethanol also tend to increase with the number of ribosomes per mRNA (Figure 3C-E). Furthermore, the ratios between the polysomal and monosomal levels of yeast RPs correlate to the corresponding ratios for their mouse orthologs (Figure 3G; p-value < 0.03), suggesting that the RP-stoichiometry differences between monosomes and polysomes are conserved across yeast and mouse.

Many yeast RPs are represented by two highly-homologous paralogs, and we explored whether the exchange among paralogs (one paralog substituting for the other) can account for the measured differential stoichiometry in Figure 3E. The levels of paralogs localized on the surface of the ribosome, such as Rpl17a and Rpl17b, are positively correlated and thus inconsistent with paralog–exchange across the analyzed ribosomes (Figure 3E). In contrast, RPs embedded deep in the core of the ribosomes either remain constant (the estimated fluctuations of their levels are within errorbars) or their paralogs exchange (e.g., the levels of Rpl37a and Rpl37b are anticorrelated; see Figure 3E), indicating that each ribosome has a copy of Rpl37. In general, the RPs whose levels differ the most among the different fractions are located on the surface of the yeast ribosomes, as can be seen from their 3D colorcoded rendition in the Supporting PDB files and Movie 1.

### RP enrichment in polysomes correlates to fitness

Next, we tested the differential RPs stoichiometry and its phenotypic consequences by independent fitness measurements. Our observation that the RP stoichiometry depends on the number of ribosomes bound per mRNA parallels measurements of higher translational activity of polysomes compared to monosomes (Warner *et al*, 1963; Goodman and Rich, 1963); some studies have even reported that the translational activity per ribosome increases with the number of ribosomes bound per mRNA (Noll *et al*, 1963; Wettstein *et al*, 1963), but this finding has not been widely reproduced. We therefore hypothesized that genetic deletions of RPs enriched in the more active ribosomes – as compared to RPs enriched in less active ribosomes – may result in a larger decrease of the translation rate and thus lower fitness. To test this hypothesis, we computed the correlation (Figure 4A) between the fitness of yeast strains with single RP–gene deletions (Qian *et al*, 2012) and the corresponding relative RP levels measured in the tetra-ribosomal fraction (4 ribosomes per mRNA). Consistent with our hypothesis, the fitness of strains lacking RP–genes is inversely proportional to the relative levels of the corresponding RPs in the tetra-ribosomes (Figure 4A). Extending this correlation analysis to the RP–levels in all sucrose fractions shown in Figure 3E results in a correlation pattern (Figure 4B) that further supports our hypothesis by showing the opposite dependence for fractions with fewer ribosomes per mRNA: the fitness of strains lacking RP–genes is proportional to the relative levels of the corresponding RPs in fractions with fewer ribosomes per mRNA (Figure 4B). This correlation pattern holds both for ethanol and for glucose carbon sources. To mitigate possible artifacts in the fitness data due to potential chromosome duplications in the deletion strains, we computed the correlations between the RP–levels and the fitness of the corresponding RP–deletion strains only for RPs without paralogs (thus unlikely to be affected by chromosome duplication) and found much higher magnitudes of the correlations (Figure 4A, B). This result suggests that the differential RP stoichiometry is not limited to paralogous RPs substituting for each other.

**Figure 4.**
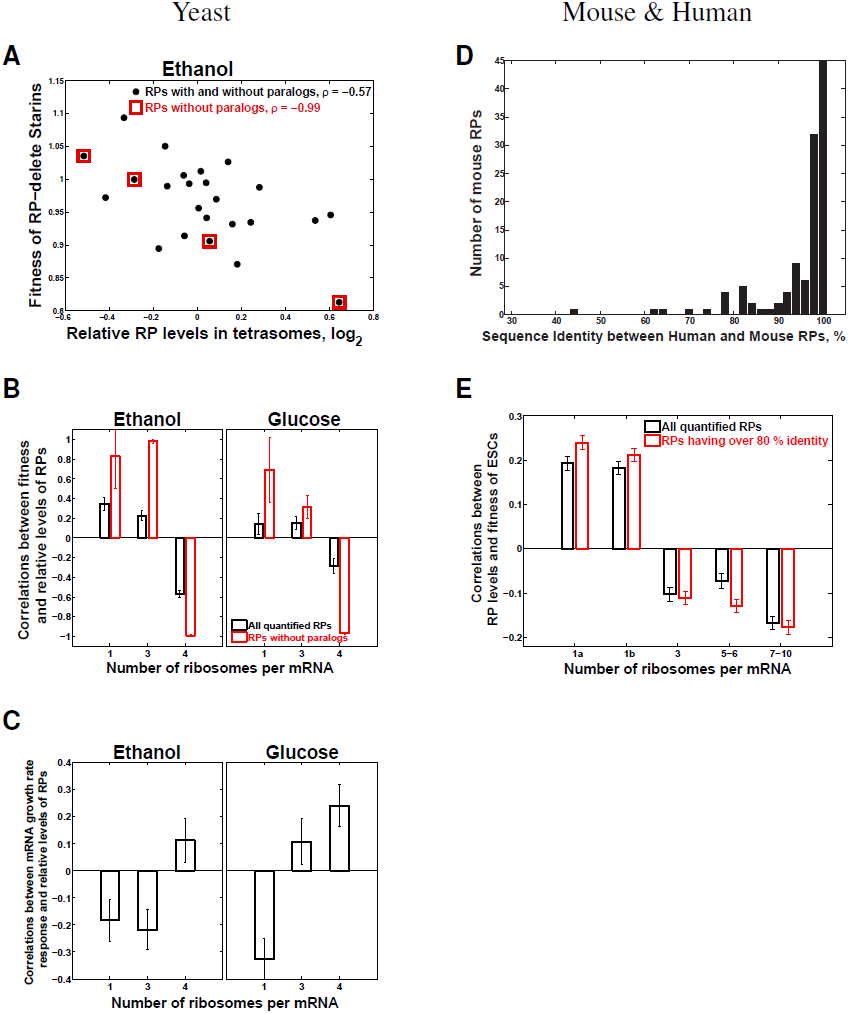
The relative levels of RPs across monosomes and polysomes correlate significantly to the fitness of yeast and mammalian cells lacking the genes encoding these RPs. (A) The fitness of RP–deleted yeast strains (Qian *et al*, 2012) is inversely proportional (p–value < 4 × 10*^−3^*) to the relative levels of the corresponding RPs in tetrasomes from yeast growing on ethanol carbon source. The RPs without paralogs are marked with red squares. (B) Extension of the analysis in panel (A) to all sucrose fractions: correlations between the relative RP levels from E and the fitnesses of strains lacking the corresponding RP genes (Qian *et al*, 2012). The correlations are shown either for all quantified RPs or only for RPs without paralogs. (C) Correlations between the relative levels of the RPs from E and the their transcriptional growth–rate responses (slopes). The growth-rate slopes were previously computed by regressing (R*^2^*> 0.87) the levels of mRNAs in glucose-limited steady-state cultures of yeast against the growth-rates of the cultures (Slavov and Botstein, 2011). (D) Distribution of sequence identity between human RPs and their closest mouse orthologs; the sequences and annotations for RPs are from Swiss–Prot. (E) Extension of the analysis for yeast in panels (A-B) to mouse: correlations between the relative levels of mouse RPs from and the fitness of human ESC lacking the corresponding human ortholog (Shalem *et al*, 2014). The correlations are shown either for all quantified RPs or only for RPs whose sequence identity between mouse and human exceeds 80 %. The correlation for monosomes is shown in replicates (1a and 1b). See also Figure S5. All error bars are SD from bootstrapping.

To further explore the functional significance of the differential RP stoichiometry, we examined whether polysome–enriched RPs are preferentially induced at higher growth–rates. We previously found that the degree of growth-rate-dependent transcriptional induction varies significantly across RPs (Brauer *et al*, 2008; Slavov and Botstein, 2011, 2013; Slavov *et al*, 2012). We quantified the growth-rate responses of RPs by regressing their mRNA levels on growth-rates and computing growth rate-slopes. The magnitudes of RP growth–rate slopes range from positive (mRNA levels increase with increasing growth rate) to negative (mRNA levels decrease with increasing growth rate), see Figure S5. Analogously to our fitness analysis (Figure 4A), we correlated the growth-rate slopes to the relative RP levels from Figure 3E. Consistent with our hypothesis, the correlation pattern (Figure 4C) indicates that the higher the growth–rate slope of a RP, the higher its enrichment in sucrose-fractions corresponding to increasing numbers of ribosomes per mRNA.

We extended our fitness analysis from yeast to mouse using the published depletion data from CRISPR knockouts in human ESC (Shalem *et al*, 2014); see Supplemental Information.

We used BLAST to identify the closest mouse orthologs of each human RP with depletion data (Figure 4D), and correlated the fitness of human ESC lacking the human RP orthologs to the RP levels across sucrose fractions that we measured, Figure 2. The correlation pattern (Figure 4E) is similar to the one in yeast (Figure 4A-C) and highly significant, FDR < 0.1%. This pattern indicates that the fitness of ESC lacking RP-genes is directly proportional to the relative RP levels in monosomes and inversely proportional to the relative RP levels in polysomes. The magnitude of this inverse proportionality increases with the number of ribosomes per mRNA (Figure 4E), consistent with our hypothesis. The fact that the fitness of human ESC lacking RPs correlates significantly to the levels of the corresponding mouse orthologous RPs suggests that the differential RP stoichiometry and its biological functions are likely conserved across mouse and human. The magnitude of this correlation increases when the correlation is computed based only on the orthologs whose sequences are over 80% identical between mouse and human (Figure 4E), providing further evidence for the conserved fitness consequences of the altered RP stoichiometry.

## Discussion

For decades, the ribosome has been considered the preeminent example of a large RNA–protein complex with a fixed stoichiometry among the constituent core RPs (Warner, 1999; Ben-Shem *et al*, 2011). However, the direct and precise measurements of RP–levels required to support this view have been very challenging. Prior to our work, the most direct and precise quantification of RP stoichiometry that we know of is based on measuring the radioactivity from RPs labeled with *^14^*C or *^3^*H and separated on 2D–gels. Some of these studies (Weber, 1972; Westermann *et al*, 1976) achieved very high precision (standard error < 10 %) and reported over 2–fold deviation from 1: 1 stoichiometry for multiple RPs. Other studies of prokaryotic ribosomes (Hardy, 1975) achieved lower precision, and the deviation from 1: 1 stoichiometry was within the experimental error of the measurements. The results reported in ref. (Weber, 1972; Westermann *et al*, 1976; Hardy, 1975) are all consistent with our findings, albeit our measurements are limited to eukaryotic ribosomes. This prior work and our measurements reflect population– averages across a heterogeneous pool of ribosomes and thus likely underestimate the magnitude of the variability among RP stoichiometries.

A simple mechanism that may account for our observations is that the rates of translation initiation and elongation depend on the RP composition. Ribosomes whose RP composition corresponds to higher ratios between the initiation and the elongation rates are likely to be found in fractions with multiple ribosomes per mRNA. Conversely, ribosomes whose RP composition corresponds to lower ratios between the initiation and the elongation rates are likely to be found in fractions with fewer ribosomes per mRNA. Indeed, increased growth–rate on glucose carbon-source that we find associated with altered RP stoichiometry has been previously reported to be associated with faster elongation rates (Bonven and Gulløv, 1979; Young and Bremer, 1976).

However, velocity sedimentation in sucrose gradients is unlikely to perfectly separate ribosomes based on their RP composition. For example, short mRNAs and the ribosomes translating them can be found only in the fractions containing few ribosomes per mRNA regardless of the efficiency of translation and the RP-composition of the ribosomes (Arava *et al*, 2003). Similarly, even the most highly translated mRNA that is likely to be translated by polysome-type ribosomes will go through a stage when only a single ribosome is loaded and thus will be found in the monosomal fraction. Other factors may also contribute to the mixing of different ribosomes in each sucrose fraction, including variation in the mRNA length, any degree of ribosome run-off, and mRNA shearing during sample handling, if any. None of these factors, however, is likely to artifactually contribute to the differential RP stoichiometry that we observe. Rather, the presence of ribosomes with different RP compositions in the same sucrose fraction would average out and decrease the differences, resulting in underestimation of the RP variability.

The conserved difference between monosomal and polysomal ribosomes (Figure 3G) raises the question about the activity of monosomes, especially given the lower estimates for their translational activity (Warner *et al*, 1963; Wettstein *et al*, 1963). The RP levels in Figure 3E indicate that the RP composition of trisomes in ethanol is more similar to the composition of monosomes than to tetrasomes. This observation shows that monosomes may have similar RP composition to polysomes, suggesting that the RP composition of monosomes is not necessarily indicative of a non–functional state.

The correlations between RP–composition and fitness can be explained by the expectation that the higher the translational activity of a ribosome, the higher the fitness cost of its perturbation in rapidly growing stem cells. The key factor required for this expectation is the differential RP stoichiometry that we measured. The differential RP stoichiometry in the absence of external perturbations suggests that cells use it as a regulatory mechanism of protein synthesis. One such example might be the preferential transcriptional induction of polysome-enriched RPs at higher growth rates (Figure 4C).

Variable mammalian RPs, such as Rps4x, Rps14, Rps20, Rpl5, Rpl10, and Rpl27, directly bind mRNAs (Castello *et al*, 2012; Kwon *et al*, 2013), and this binding might mediate translational regulation as previously suggested (Mauro and Edelman, 2002; Landry *et al*, 2009; Mazumder *et al*, 2003). Furthermore, deletions or overexpressions of many of the variable RPs (Figure 1B) have well characterized phenotypes both in development and in cancer. For example, the knockdown or haploinsufficiency of the polysomally enriched Rps19 (Figure 1B) causes Diamond Blackfan anemia by selectively affecting the synthesis of some proteins but not of others (Horos *et al*, 2012). Interestingly, our data indicate that RPs that are frequently mutated in cancers, such as Rpl5 and Rpl10 (De Keersmaecker *et al*, 2013; Lawrence *et al*, 2014), are enriched in the monosomes (Figure 1A and Figure 2). Conversely, RPs whose (over)– expression promotes cancer, such as Rpl30, Rps20, and Rpl39 (De Bortoli *et al*, 2006; Dave *et al*, 2014), are enriched in the polysomes (Figure 1B and Figure 2). One interpretation, among others, of these data is that loss of function of monosomally–enriched RPs or overexpression of polysomally–enriched RPs might promote protein synthesis and cancer cell growth.

## Experimental Procedures

All yeast experiments used a prototrophic diploid strain (DBY12007) with a S288c background and wild type HAP1 alleles (Slavov and Botstein, 2011). We grew our cultures in a bioreactor (LAMBDA Laboratory Instruments) using minimal media with the composition of yeast nitrogen base (YNB) and supplemented with 2g/L D-glucose.

Mouse embryonic stem cells (E14 10*^th^* passage) were grown as adherent cultures in 10 cm plates with 10 ml DMEM/F12 media supplemented with 10 % knockout serum replacement, nonessential amino acids (NEAA supplement), 0.1 mM β–mercapto–ethanol, 1 % penicillin and streptomycin, leukemia inhibitory factor (LIF; 1,000 U LIF/ml), and 2i (GSK3β and Mek 1/2 inhibitors).

Both yeast and mouse embryonic stem cells were lysed by vortexing for 10 min with glass beads in cold PLB buffer. The crude extracts obtained from this lysis procedure were clarified by centrifugation. The resulting supernatants were applied to linear 11 ml sucrose gradients (10 % − 50 %) and spun at 35,000 rpm in a Beckman SW41 rotor either for 3 hours (for yeast samples) or for 2.5 hours (for mouse samples). Twelve fractions from each sample were collected using a Gradient Station. More details are available in the Supplemental Information.

## Data availability

The raw MS data have been deposited in MassIVE (ID: MSV000079280) and in the ProteomeXchange (ID: PXD002816) Supplementary materials, data, and 3D ribosomal structures color–coded according to the RP levels from Figure 3E can be found at: http://alum.mit.edu/www/nslavov/Ribosome_Data/

## Acknowledgments

We thank J. Cate and N. Lintner for helping us colorcode the variability of RPs on the 3D structure of the yeast ribosomes, P. Vaidyanathan for help with the sucrose gradients, R. Robertson for technical assistance, and M. Jovanovic, Y. Katz, S. Kryazhimskiy, W. Gilbert, P. Vaidyanathan, G. Frenkel, D. Mooijman, J. Alvarez, D. Botstein, and A. Murray for discussions and constructive comments. This work was funded by a grant from the National Institutes of Health to A.v.O. (R01-GM068957) and a SPARC grant to N.S.

## Author Contributions

Conceptualization, N.S.; Methodology, N.S.; Investigation, N.S., S.S., and B.B.; Writing Original Draft, N.S.; Writing Review & Editing, N.S., S.S., and A.v.O.; Funding Acquisition, A.v.O and N.S.; Resources, A.v.O, E.A., and N.S.; Supervision, N.S.

## References

Arava Y, Wang Y, Storey JD, Liu CL, Brown PO, Herschlag D (2003) Genome-wide analysis of mRNA translation profiles in *Saccharomyces cerevisiae*. Proceedings of the National Academy of Sciences 100: 3889–3894

Ashe MP, Susan K, Sachs AB (2000) Glucose depletion rapidly inhibits translation initiation in yeast. Molecular Biology of the Cell 11: 833–848

Ben-Shem A, de Loubresse NG, Melnikov S, Jenner L, Yusupova G, Yusupov M (2011) The structure of the eukaryotic ribosome at 3.0 Å resolution. Science 334: 1524–1529

Bonven B, Gulløv K (1979) Peptide chain elongation rate and ribosomal activity in *Saccharomyces cerevisiae* as a function of the growth rate. Molecular and General Genetics MGG 170: 225–230

Brauer MJ, Huttenhower C, Airoldi EM, Rosenstein R, Matese JC, Gresham D, Boer VM, Troyanskaya OG, Botstein D (2008) Coordination of Growth Rate, Cell Cycle, Stress Response, and Metabolic Activity in Yeast. Mol Biol Cell 19: 352–367

Castelli LM, Lui J, Campbell SG, Rowe W, Zeef LA, Holmes LE, Hoyle NP, Bone J, Selley JN, Sims PF, et al (2011) Glucose depletion inhibits translation initiation via eIF4A loss and subsequent 48S preinitiation complex accumulation, while the pentose phosphate pathway is coordinately up-regulated. Molecular biology of the cell 22: 3379–3393

Castello A, Fischer B, Eichelbaum K, Horos R, Beckmann BM, Strein C, Davey NE, Humphreys DT, Preiss T, Steinmetz LM, et al (2012) Insights into RNA biology from an atlas of mammalian mRNA–binding proteins. Cell 149: 1393–1406

Chen SS, Williamson JR (2013) Characterization of the Ribosome Biogenesis Landscape in *E. coli* Using Quantitative Mass Spectrometry. Journal of molecular biology 425: 767–779

Dave B, Granados-Principal S, Zhu R, Benz S, Rabizadeh S, Soon-Shiong P, Yu KD, Shao Z, Li X, Gilcrease M, et al (2014) Targeting RPL39 and MLF2 reduces tumor initiation and metastasis in breast cancer by inhibiting nitric oxide synthase signaling. Proceedings of the National Academy of Sciences: 201320769

De Bortoli M, Castellino RC, Lu XY, Deyo J, Sturla LM, Adesina AM, Perlaky L, Pomeroy SL, Lau CC, Man TK, et al (2006) Medulloblastoma outcome is adversely associated with overexpression of EEF1D, RPL30, and RPS20 on the long arm of chromosome 8. BMC cancer 6: 223

De Keersmaecker K, Atak ZK, Li N, Vicente C, Patchett S, Girardi T, Gianfelici V, Geerdens E, Clappier E, Porcu M, et al (2013) Exome sequencing identifies mutation in CNOT3 and ribosomal genes RPL5 and RPL10 in T-cell acute lymphoblastic leukemia. Nature genetics 45: 186–190

Fabian MR, Sonenberg N (2012) The mechanics of miRNA-mediated gene silencing: a look under the hood of miRISC. Nature structural molecular biology 19: 586–593

Fortier S, MacRae T, Bilodeau M, Sargeant T, Sauvageau G (2015) Haploinsufficiency screen highlights two distinct groups of ribosomal protein genes essential for embryonic stem cell fate. Proceedings of the National Academy of Sciences 112: 2127–2132

Galkin O, Bentley AA, Gupta S, Compton BA, Mazumder B, Kinzy TG, Merrick WC, Hatzoglou M, Pestova TV, Hellen CU, et al (2007) Roles of the negatively charged N-terminal extension of *Saccharomyces cerevisiae* ribosomal protein S5 revealed by characterization of a yeast strain containing human ribosomal protein S5. RNA 13: 2116–2128

Gilbert WV (2011) Functional specialization of ribosomes? Trends in biochemical sciences 36: 127–132

Goodman HM, Rich A (1963) Mechanism of polyribosome action during protein synthesis. Nature 199: 318–322

Gupta V, Warner JR (2014) Ribosome-omics of the human ribosome. rna 20: 1004–1013

Hardy SJ (1975) The stoichiometry of the ribosomal proteins of *Escherichia coli*. Molecular and General Genetics MGG 140: 253–274

Hendrickson DG, Hogan DJ, McCullough HL, Myers JW, Herschlag D, Ferrell JE, Brown PO (2009) Concordant regulation of translation and mRNA abundance for hundreds of targets of a human microRNA. PLoS biology 7: e1000238

Horos R, IJspeert H, Pospisilova D, Sendtner R, Andrieu-Soler C, Taskesen E, Nieradka A, Cmejla R, Sendtner M, Touw IP, et al (2012) Ribosomal deficiencies in Diamond–Blackfan anemia impair translation of transcripts essential for differentiation of murine and human erythroblasts. Blood 119: 262–272

Jenner L, Melnikov S, de Loubresse NG, Ben-Shem A, Iskakova M, Urzhumtsev A, Meskauskas A, Dinman J, Yusupova G, Yusupov M (2012) Crystal structure of the 80S yeast ribosome. Current opinion in structural biology 22: 759–767

Jovanovic M, Rooney MS, Mertins P, Przybylski D, Chevrier N, Satija R, Rodriguez EH, Fields AP, Schwartz S, Raychowdhury R, et al (2015) Dynamic profiling of the protein life cycle in response to pathogens. Science 347: 1259038

Komili S, Farny NG, Roth FP, Silver PA (2007) Functional specificity among ribosomal proteins regulates gene expression. Cell 131: 557–571

Kondrashov N, Pusic A, Stumpf CR, Shimizu K, Hsieh AC, Xue S, Ishijima J, Shiroishi T, Barna M (2011) Ribosome-mediated specificity in Hox mRNA translation and vertebrate tissue patterning. Cell 145: 383–397

Kwon SC, Yi H, Eichelbaum K, Föhr S, Fischer B, You KT, Castello A, Krijgsveld J, Hentze MW, Kim VN (2013) The RNA-binding protein repertoire of embryonic stem cells. Nature structural molecular biology

Landry DM, Hertz MI, Thompson SR (2009) RPS25 is essential for translation initiation by the Dicistroviridae and hepatitis C viral IRESs. Genes development 23: 2753–2764

Lawrence MS, Stojanov P, Mermel CH, Robinson JT, Garraway LA, Golub TR, Meyerson M, Gabriel SB, Lander ES, Getz G (2014) Discovery and saturation analysis of cancer genes across 21 tumour types. Nature 505: 495–501

Lee ASY, Burdeinick-Kerr R, Whelan SP (2013) A ribosome-specialized translation initiation pathway is required for cap-dependent translation of vesicular stomatitis virus mRNAs. Proceedings of the National Academy of Sciences 110: 324–329

Mauro VP, Edelman GM (2002) The ribosome filter hypothesis. Proceedings of the National Academy of Sciences 99: 12031–12036

Mazumder B, Sampath P, Seshadri V, Maitra RK, DiCorleto PE, Fox PL (2003) Regulated release of L13a from the 60S ribosomal subunit as a mechanism of transcript-specific translational control. Cell 115: 187–198

Mulder AM, Yoshioka C, Beck AH, Bunner AE, Milligan RA, Potter CS, Carragher B, Williamson JR (2010) Visualizing ribosome biogenesis: parallel assembly pathways for the 30S subunit. Science 330: 673–677

Noll H, Staehelin T, Wettstein F (1963) Ribosomal aggregates engaged in protein synthesis: ergosome breakdown and messenger ribonucleic acid transport. Nature 198: 632–638

O’Leary MN, Schreiber KH, Zhang Y, Duc ACE, Rao S, Hale JS, Academia EC, Shah SR, Morton JF, Holstein CA, et al (2013) The ribosomal protein Rpl22 controls ribosome composition by directly repressing expression of its own paralog, Rpl22l1. PLoS genetics 9: e1003708

Ori A, Banterle N, Iskar M, Andrés-Pons A, Escher C, Khanh Bui H, Sparks L, Solis-Mezarino V, Rinner O, Bork P, et al (2013) Cell type-specific nuclear pores: a case in point for context-dependent stoichiometry of molecular machines. Molecular systems biology 9

Parenteau J, Durand M, Morin G, Gagnon J, Lucier JF, Wellinger RJ, Chabot B, Abou Elela S (2011) Introns within ribosomal protein genes regulate the production and function of yeast ribosomes. Cell 147: 320–331

Qian W, Ma D, Xiao C, Wang Z, Zhang J (2012) The genomic landscape and evolutionary resolution of antagonistic pleiotropy in yeast. Cell Reports 2: 1399–1410

Ramagopal S (1990) Induction of cell-specific ribosomal proteins in aggregation competent nonmorphogenetic *Dictyostelium discoideum*. Biochemistry and Cell Biology 68: 1281–1287

Ramagopal S, Ennis HL (1981) Regulation of synthesis of cell-specific ribosomal proteins during differentiation of *Dictyostelium discoideum*. Proceedings of the National Academy of Sciences 78: 3083–3087

Reschke M, Clohessy JG, Seitzer N, Goldstein DP, Breitkopf SB, Schmolze DB, Ala U, Asara JM, Beck AH, Pandolfi PP (2013) Characterization and Analysis of the Composition and Dynamics of the Mammalian Riboproteome. Cell Reports 4: 1276–1287

Sampath P, Pritchard DK, Pabon L, Reinecke H, Schwartz SM, Morris DR, Murry CE (2008) A hierarchical network controls protein translation during murine embryonic stem cell self-renewal and differentiation. Cell stem cell 2: 448–460

Shalem O, Sanjana NE, Hartenian E, Shi X, Scott DA, Mikkelsen TS, Heckl D, Ebert BL, Root DE, Doench JG, et al (2014) Genome-scale CRISPR-Cas9 knockout screening in human cells. Science 343: 84–87

Slavov N, Airoldi EM, van Oudenaarden A, Botstein D (2012) A conserved cell growth cycle can account for the environmental stress responses of divergent eukaryotes. Molecular Biology of the Cell 23: 1986–1997

Slavov N, Botstein D (2011) Coupling among growth rate response, metabolic cycle, and cell division cycle in yeast. Molecular Biology of the Cell 22: 1997–2009

Slavov N, Botstein D (2013) Decoupling Nutrient Signaling from Growth Rate Causes Aerobic Glycolysis and Deregulation of Cell-Size and Gene Expression. Molecular Biology of the Cell 24: 157–168

Slavov N, Budnik B, Schwab D, Airoldi E, van Oudenaarden A (2014) Constant Growth Rate Can Be Supported by Decreasing Energy Flux and Increasing Aerobic Glycolysis. Cell Reports 7: 705–714

Slavov N, Dawson KA (2009) Correlation signature of the macroscopic states of the gene regulatory network in cancer. Proceedings of the National Academy of Sciences 106: 4079–4084

Vaidyanathan PP, Zinshteyn B, Thompson MK, Gilbert WV (2014) Protein kinase A regulates gene-specific translational adaptation in differentiating yeast. RNA: 10.1261/rna.044552.114

Wang X, Chen CF, Baker PR, Chen Pl, Kaiser P, Huang L (2007) Mass spectrometric characterization of the affinity-purified human 26S proteasome complex. Biochemistry 46: 3553–3565

Warner JR (1999) The economics of ribosome biosynthesis in yeast. Trends in biochemical sciences 24: 437–440

Warner JR, Knopf PM, Rich A (1963) A multiple ribosomal structure in protein synthesis. Proceedings of the National Academy of Sciences of the United States of America 49: 122

Warner JR, McIntosh KB (2009) How common are extraribosomal functions of ribosomal proteins? Molecular cell 34: 3–11

Weber HJ (1972) Stoichiometric measurements of 30S and 50S ribosomal proteins from *Escherichia coli*. Molecular and General Genetics MGG 119: 233–248

Westermann P, Heumann W, Bielka H (1976) On the stoichiometry of proteins in the small ribosomal subunit of hepatoma ascites cells. FEBS letters 62: 132–135

Wettstein F, Staehelin T, Noll H (1963) Ribosomal aggregate engaged in protein synthesis: characterization of the ergosome. Nature 197: 430–435

Wool IG (1996) Extraribosomal functions of ribosomal proteins. Trends in biochemical sciences 21: 164–165

Xue S, Barna M (2012) Specialized ribosomes: a new frontier in gene regulation and organismal biology. Nature Reviews Molecular Cell Biology 13: 355–369

Young R, Bremer H (1976) Polypeptide-chain-elongation rate in *Escherichia coli* B/r as a function of growth rate. Biochem J 160: 185–194

